# A temporal gap in sensory streams amplifies the influence of subsequent input on decision making

**DOI:** 10.1101/2024.07.17.603868

**Authors:** Alejandro Sospedra, Santiago Canals, Encarni Marcos

## Abstract

Decision making involves evaluating options and predicting their likely outcomes. Traditional laboratory studies of decision making often employ tasks involving the discrimination of perceptual evidence, where sensory information is constant and presented continuously. However, during natural behavior, decision making usually involves intermittent information streams, punctuated by periods with no input. To investigate decision making under such conditions, we designed a perceptual task where participants observed tokens sequentially jumping from a central circle to one of two peripheral targets, disappearing shortly after. Participants were required to report which target they believed would have received most tokens by the trial’s end. Half of the trials included a temporal gap, during which no information was displayed. To better understand decision-making dynamics, we introduced specific patterns of token jumps. We found that participants made choices with less available information and disproportionally weighted the information presented immediately after the gap more heavily than they did when no gap was present. Traditional computational models, which assume uniform or gradual decay or increase of weighting of information over time, could not account for this effect. A control task with randomly presented information further confirmed that the disproportionate weighting of post-gap information is a robust feature of the decision-making process. These findings highlight the importance of studying decision making in environments with intermittent information and temporal gaps, where the timing and structure of inputs critically shape behavior.

**Significance statement:** Decision making in real-world environments often involves intermittent information, where key evidence might be separated by temporal gaps. Our study demonstrate that people disproportionately rely on information received immediately after such gaps, weighting the same information more heavily than when no gap is present. Traditional decision-making models do not account for this effect, as they lack mechanisms to capture how temporal disruptions shape evidence weighting. By understanding how gaps influence decision making, this research provides insights into how people adapt their decisions in fluctuating environments, with implications for fields ranging from neuroscience to artificial intelligence.

## Introduction

Perceptual decision making is generally considered to be a deliberative process, where individuals evaluate sensory evidence over time to make informed choices (Smith and Ratcliff, 2004; Gold and Shadlen, 2007). In most laboratory studies, sensory information is presented consistently, with a stable amount of evidence favoring a particular choice. While such paradigms have been pivotal in understanding decision-making dynamics, they often fail to capture the complexity of real-world decision making, where sensory evidence is frequently intermittent, fluctuating, and punctuated by periods where no new information is available. Some experimental designs have attempted to address this limitation by incorporating dynamic information, where the net evidence fluctuates over time (Cisek et al., 2009; Thura and Cisek, 2014; Winkel et al., 2014; Holmes et al., 2016; Ferrucci et al., 2021; Trueblood et al., 2021). However, even these studies often assume that information remains continuously available, neglecting the unique challenges posed by intermittent evidence. In natural environments, individuals must adapt to interruptions in sensory input, requiring distinct strategies that may fundamentally alter how evidence is weighted.

Traditionally, decision-making tasks typically present continuous sensory evidence, requiring animals or humans to report their decisions at the end of such streams of information (Ratcliff, 1978; Shadlen and Newsome, 2001; Roitman and Shadlen, 2002; Mazurek et al., 2003; Kiani et al., 2008; Kiani et al., 2013). In these paradigms, where the net sensory evidence consistently favors one choice, perceptual decisions appear unaffected by periods without information, leading to the proposal that a neural mechanism “freezes” the decision-making process during interruptions (Kiani et al., 2013; Waskom and Kiani, 2018; Tohidi-Moghaddam et al., 2019; Azizi and Ebrahimpour, 2023). This stability is often attributed to sustained neuronal activity that remains robust to external forces (Marcos et al., 2019). However, these findings might be confounded by different factors. For instance, the continuous and consistent nature of the evidence before and after temporal gaps allows participants to rely on the most recent evidence, without the need to maintain or integrate all prior information. Additionally, focusing solely on accuracy limits the interpretation of the decision process, as it overlooks critical variables such as decision times and the temporal structure of evidence (Kiani et al., 2013). Recent studies have emphasized the importance of incorporating additional variables, such as fluctuations in overall sensory evidence, decision times and the temporal placement of information, to better understand decision making (Kiani et al., 2008; Cisek et al., 2009; Thura et al., 2012; Brunton et al., 2013; Hanks et al., 2015; Carland et al., 2016; Piet et al., 2018; Ferrucci et al., 2021; Trueblood et al., 2021).

Here, we directly examine how temporal interruptions –combined with fluctuating sensory evidence – affect both the timing of decisions and the weighting of evidence. Using a behavioral task that introduces a temporal gap within a stream of dynamically changing evidence, we aim to provide a deeper understanding of decision making under conditions of intermittent sensory input. By combining this task with computational modeling, we explore how such interruptions influence the decision-making dynamics and assess whether traditional models can adequately explain these effects. Our task required participants to track perceptual information, which changed in discrete steps, and make decisions at any point during the trial. This design not only allowed us to measure decision times but also to precisely assess the specific influence of each individual piece of information on decisions. By examining how interruptions affect the processing of dynamic evidence, our study represents a step toward bridging the gap between controlled laboratory paradigms and the complexities of real-world decision making.

## Results

The experimental tasks were based on the tokens task (Cisek et al., 2009; Ferrucci et al., 2021) (see Materials and Methods). Briefly, fifteen tokens appeared within a central circle on the screen and sequentially jumped to one of two peripheral circles (targets), disappearing before the next token made its jump (Fig 1A). In half of the trials, the sequence of tokens occurred without interruption (no-pause trials; Fig 1B-top), while in the other half, a temporal gap of 300 ms was introduced during the sequence (pause trials; Fig 1B-bottom). Participants were instructed to predict, at any time, which of the two targets would receive more tokens by the end of the trial and report their choice by placing the mouse cursor on the chosen target. Two experimental tasks were designed: in the structured task, specific jump patterns were introduced to create well-defined temporal sequences (Fig 1C), while in the non-structured task, which served as control, no predefined patterns were applied to the tokens jumps. This control task ensured that any observed effects in the structured task were specifically attributed to the temporal dynamics of the stimulus rather than being driven by implicit salience effects due to the introduced token sequences.

**Figure 1.**
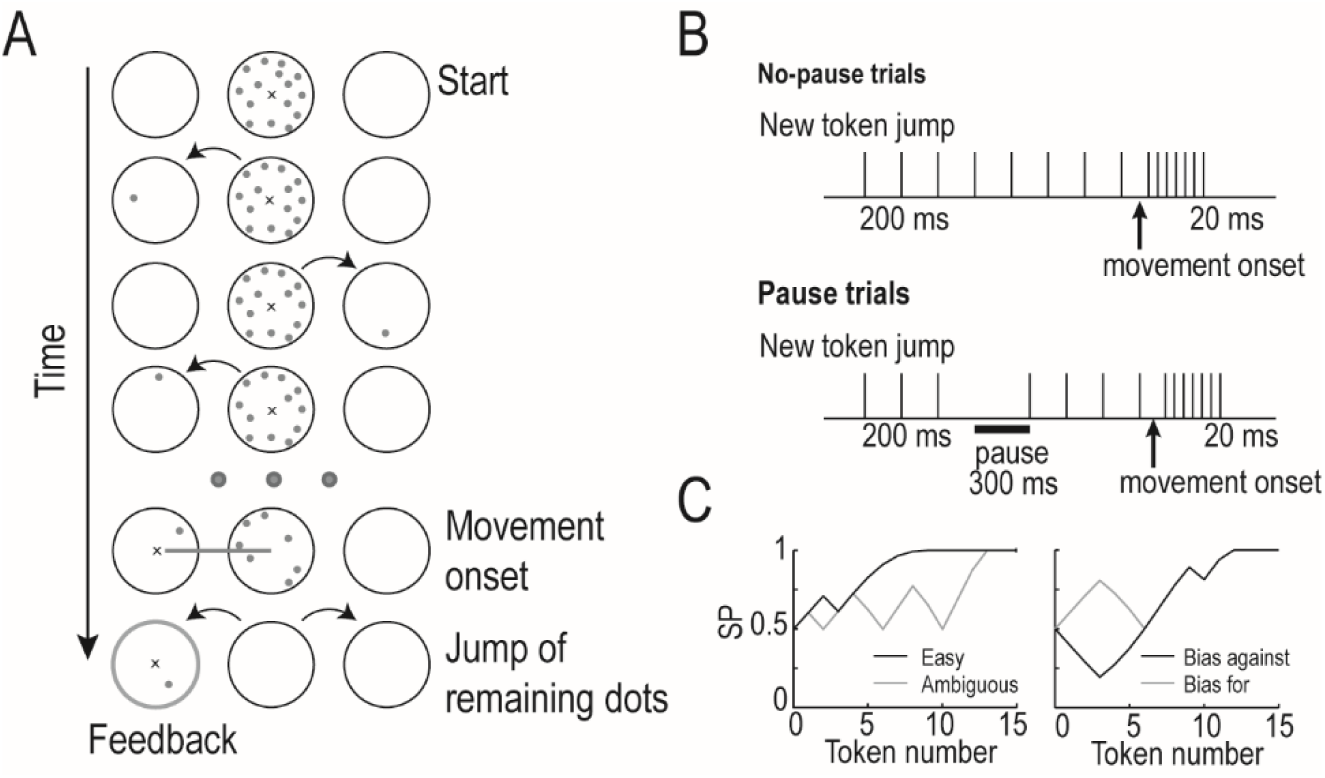
Experimental design. **(A)** Temporal sequence of task events. At the beginning of each trial, fifteen tokens are presented on a central circle on the screen. When the participant moves the mouse cursor within the central circle, the tokens start to sequentially jump (every 200 ms) to one of the two peripheral circles (targets), disappearing shortly after. Participants are asked to select the peripheral target that they think will accumulate the majority of the tokens by the end of the trial. To make their choice, they have to move the cursor within the selected target. **(B)** Sequential order of tokens’ jumps for no-pause (*top*) and pause trials (*bottom*). **(C)** Success probability (SP) for specific trial’s profiles: easy and ambiguous trials (*left*), bias-for and bias- against trials (*right*).

Overall, participants performed the structured task with an accuracy well above chance (mean accuracy: 78 ± 1 %), indicating a strong ability to discriminate the sensory evidence. To assess how behavior was modulated by the trial profile, we analyzed decision times (DTs), success probabilities (SPs) and accuracy across the various trial profiles. Specifically, we compared performance between easy and ambiguous trials to determine whether the overall difficulty of the trial influenced behavior (Fig 1C-left). Additionally, we contrasted bias-for and bias-against trials, focusing only on those trials where responses were made after the 7^th^ token. This time point was chosen because it marks the moment when the two trial profiles converge in terms of SPs for the reminder of the trial (Fig 1C-right). Any differences in behavior for decisions made after this token would indicate the influence of prior information on choices. Given that information is not always available, any modulation of behavior across trials profiles suggests the need for working memory to track and maintain relevant information (Ferrucci et al., 2021).

DTs varied depending on the trial profile, with shorter DTs observed for easy trials compared to ambiguous trials and for bias-against compared to bias-for trials (Fig 2-left; paired-samples Wilcoxon signed-rank test, z = 4.01 and z = 3.49, respectively; false discovery rate (FDR)-corrected for multiple comparisons p < 0.001). SPs at the time of decision were also significantly different across conditions (Fig 2-middle; paired-samples Wilcoxon signed-rank test, z = 3.39 - 4.01, FDR-corrected for multiple comparisons, p < 0.001). Similarly, accuracy was significantly modulated by trial condition (Fig 2-right; paired-samples Wilcoxon signed-rank test z = 4.01 for both comparisons, FDR-corrected for multiple comparisons, p < 0.0001). Consistent with previous studies, these results indicate that decisions are influenced by the temporal structure of the information flow, leading to a modulation of behavior, and that working memory plays a role in monitoring and integrating prior information throughout the decision-making process (Cisek et al., 2009; Ferrucci et al., 2021). Next, we asked whether such modulation could also be influenced by the presence or absence of the temporal gap.

**Figure 2.**
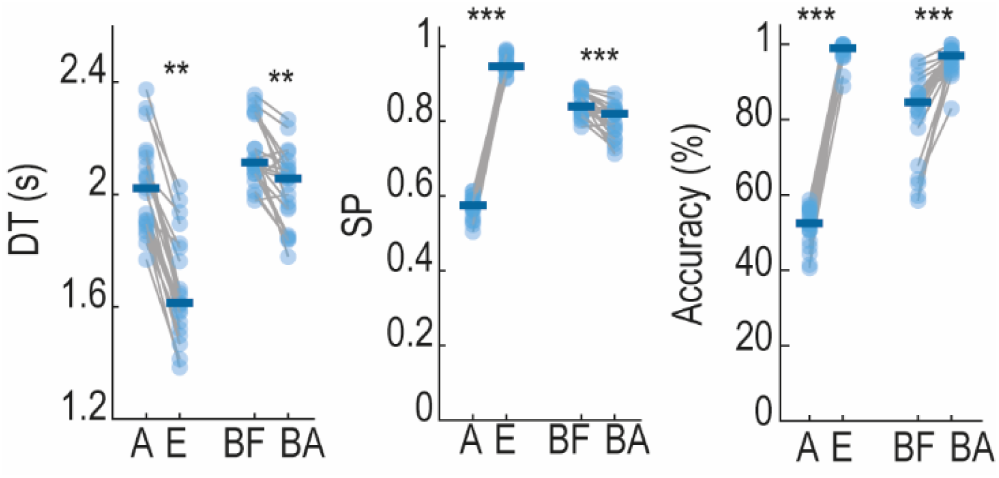
**Behavior of subjects for different trial types**. DT (*left*), SP (*middle*) and accuracy (*right*) for ambiguous (A), easy (E), bias-for (BF) and bias-against trials (BA) (paired Wilcoxon signed rank test, FDR-corrected for multiple comparisons, *** = p<0.0001, ** = p<0.001). Colored solid line shows the median of the population.

### Decisions are influenced by a temporal gap

To investigate the influence of a temporal gap in decisions, we analyzed only the trials where responses occurred after the 4^th^ token, ensuring that the gap was present in pause trials, which occurred between the 3^rd^ and the 4^th^ token. Participants’ responses were significantly modulated by the presence of the temporal gap. They exhibited differences in their DTs between no-pause and pause trials, showing shorter DTs in the absence of a pause compared to trials with a pause (Fig 3A-left; paired-samples Wilcoxon signed-rank test, z = 4.01, p < 0.0001). Importantly, the time difference was shorter than the duration of the pause (mean difference in DTs ± SEM: 0.173 ± 0.123 s *vs*. 0.3 s), indicating that decision making progresses during the pause. If this were not the case, the DT difference would have matched the pause duration.

**Figure 3.**
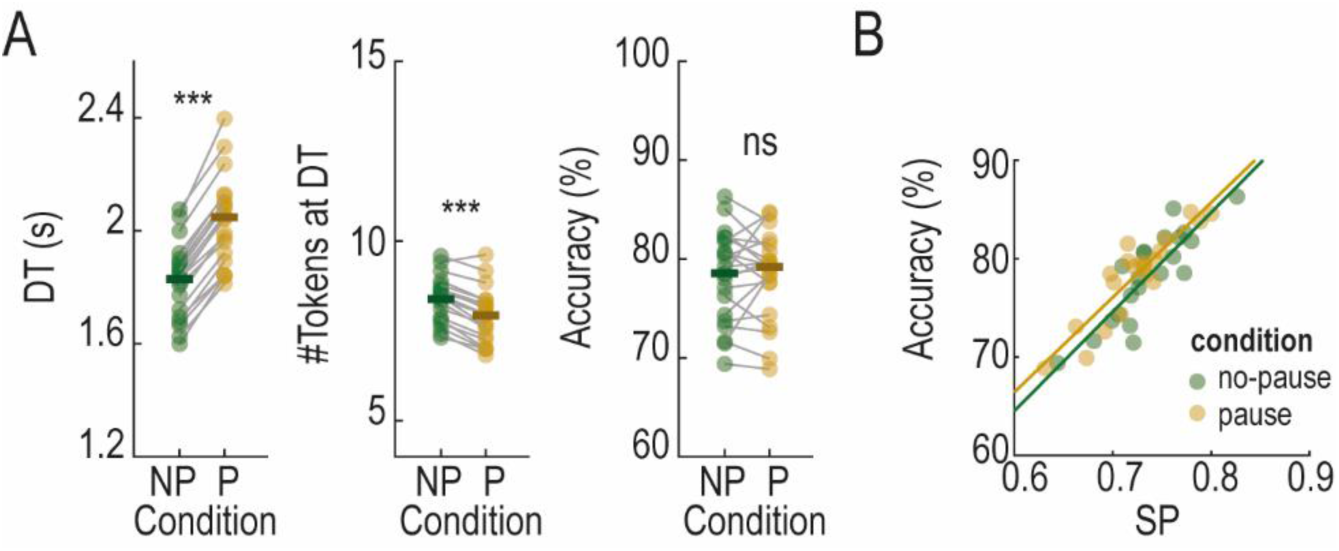
Behavior for subjects for pause and no-pause trials. **(A)** DTs (*left*), number of tokens at DTs (*middle*) and accuracy (*right*) divided into no-pause (NP) and pause (P) trials (paired Wilcoxon signed-rank test, *** = p < 0.0001, ns = non-significant). **(B)** SP Vs accuracy for all correct and incorrect trials divided into pause and no-pause conditions.

However, despite the prolonged DTs observed during pause trials, participants did not utilize more information to make decisions. Instead, decisions were made with fewer tokens when a pause was present in the trial (Fig 3A-middle; paired-samples Wilcoxon signed-rank test, z = 3.98, p < 0.0001). Consequently, choices were made with less SP for pause than no-pause trials (mean SP ± SEM: 0.80 ± 0.01 and 0.82 ± 0.01 for no-pause and pause trials, respectively; paired-samples Wilcoxon signed-rank test, z = 3.81, p < 0.001). Interestingly, such reduction in both the number of tokens required for decision making and the SP at DT did not compromise performance. Accuracy was comparable across conditions (Fig 3A-right; paired-samples Wilcoxon signed-rank test, p = 0.77), and significantly exceeded the accuracy predicted by the decrease in SP at the time of decision (paired-samples Wilcoxon signed-rank test z = 2.90, p < 0.01). This prediction was derived from the fitted relationship between SPs and accuracy (Fig 3B).

To investigate whether the presence of a pause modulates the differences observed between trial profiles, we analyzed the data by separating it into different trial types. To do this, we considered trials with decisions made after the jump of the 4^th^ token for easy and ambiguous trials, and after the jump of the 7^th^ token for bias-for and bias-against trials. Significant differences in DTs were observed between easy and ambiguous trials, as well as between bias-against and bias-for trials, in both no-pause and pause conditions (Fig 4-left; paired-samples Wilcoxon signed-rank test, z = 3.22 - 4.01, FDR-corrected for multiple comparisons, p < 0.01). Similarly, accuracy was modulated by the trial type in both no-pause and pause conditions, mirroring the trends observed in the main behavioral analysis (Fig 2-right and Fig 4-right; paired-samples Wilcoxon signed-rank test, z = 3.91 - 4.01, FDR-corrected for multiple comparisons, p < 0.0001). These findings align with the idea of a working memory process that retain information not visually available, which then degrades over the course of the trial (Ferrucci et al., 2021). Moreover, the difference in the modulation of DTs between bias-for and bias-against trials for no-pause and pause conditions was not significantly different (paired-samples Wilcoxon signed-rank test, p > 0.9), suggesting that the temporal gap does not increase the degradation of prior information. If it did, we would expect an increase in the difference between DTs for the pause condition.

**Figure 4.**
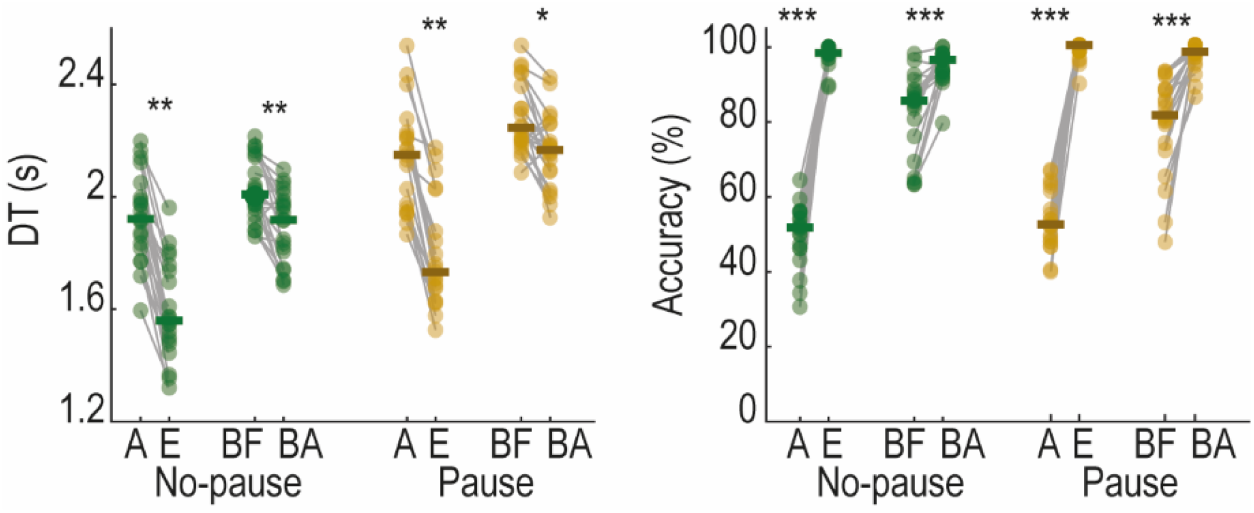
**Behavioral modulation across trial types for pause and no-pause conditions**. DTs (*left*) and accuracy across trial types (*right*) for pause and no-pause conditions. In both cases, only correct trials with decisions made after the jump of the 4th token for ambiguous (A) and easy (E) trials and after the jump of the 7th token for bias-for (BF) and bias-against trials (BA) were considered (paired-samples Wilcoxon signed-rank test, *** = p < 0.0001, *** = p < 0.001, * = p < 0.01).

Therefore, a temporal gap introduced within the course of some trials modulated DTs of choices and the perceived amount of information. Surprisingly, however, this manipulation did not affect the accuracy of choices. This raises a pivotal question: can traditional decision-making models account for these effects, or do they require a fundamental revision to explain behavior under intermittent information streams?

### Traditional decision-making models fail to fully explain behavior

We assessed existing models of decision making to determine whether they can account for the influence of temporal gaps on choices under conditions where information is incomplete and/or delayed. To explore this, we examined three well-established models from the literature: a drift diffusion model with fixed bounds (DDM-fixed), a drift diffusion model with collapsing bounds (DDM-collapsing) and an urgency-gating model (UGM). The models were selected because of their ability to explain a wide variety of behavior (Stone, 1960; Laming, 1968; Ratcliff, 1978; Usher and McClelland, 2001; Mazurek et al., 2003; Bogacz and Gurney, 2007; Hawkins et al., 2015; Ferrucci et al., 2021; Yau et al., 2021; Smith and Ratcliff, 2022). In all these cases, working memory is responsible for storing and updating past information with incoming data (Ferrucci et al., 2021). To account for the observed differences between bias-for and bias-against trials, as well as the absence of modulation of this difference in pause trials, we introduced a leakage term into the working memory component. This term caused the degradation of stored information only when new information arrived (see Materials and Methods).

To evaluate how well the computational models captured the observed behavior, we used a differential evolution algorithm to fit the experimental data. The estimated parameters for each model are shown in Table 1. Both the DDM-collapsing and UGM provided substantially better fits to the data than the DDM-fixed, as evidenced by the shapes of the real and simulated DT distributions (Fig 5A-C). This result was further confirmed by significantly better Bayesian Information Criterion (BIC) values for the DDM-collapsing and UGM compared to the DDM- fixed across subjects (paired-samples Wilcoxon signed-rank test, z = 3.53 – 3.81, FDR-corrected for multiple comparisons, p < 0.001). This underscores the critical importance of incorporating an internal signal related to the urgency to respond in order to account for behavior, highlighting that the combination of perceptual evidence with urgency is essential to understand most decisions.

**Figure 5.**
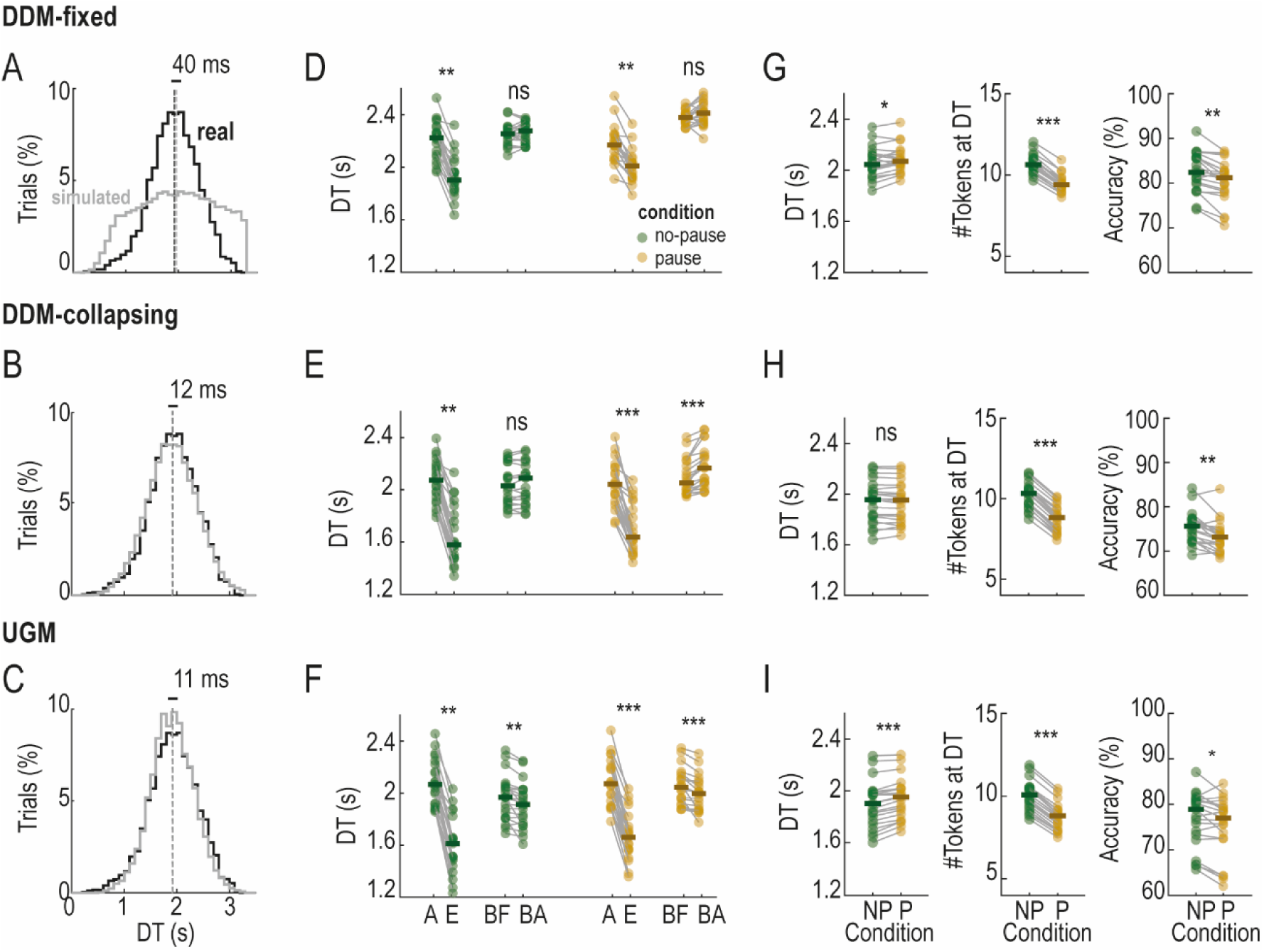
Simulated results for the best fitting parameters. (A-C) DTs distribution for real and simulated data for the DDM-fixed, DDM-collapsing and UGM, pooled across all subjects (real data) and all simulations (simulated data). **(D-E)** DTs for ambiguous (A), easy (E), bias-for and bias-against (BA) trials types across no-pause and pause conditions. **(G-I)** DTs (*left*), number of tokens at decision (*middle*) and accuracy (*right*) for no-pause (NP) and pause (P) conditions when only trials with decisions made after the 4^th^ token are considered (paired-samples Wilcoxon signed-rank test, *** = p<0.0001, ** = p<0.001, * = p<0.05; ns, non-significant). The simulated data was obtained with 5,000 trials for each model.

**Table 1.**
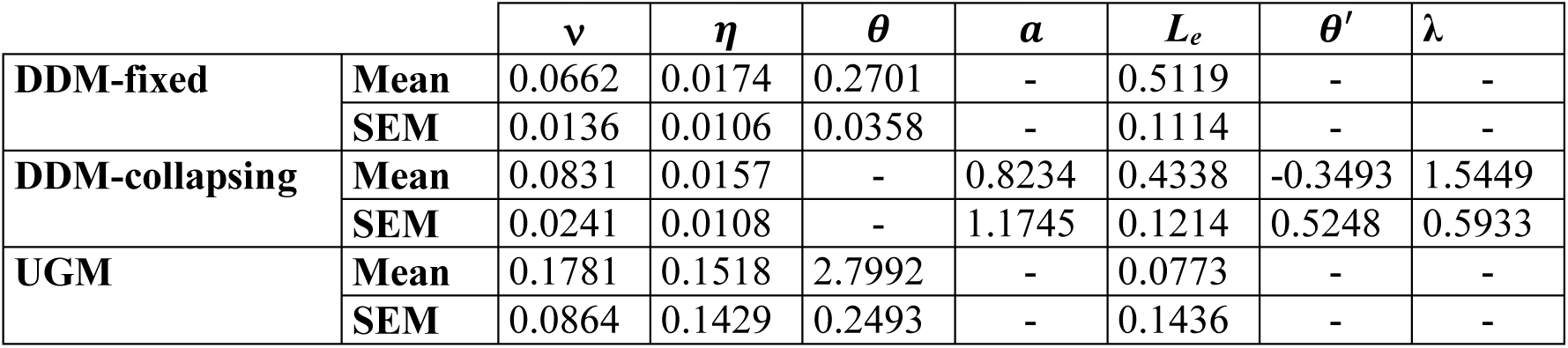
Value of the free parameters estimated for the three models. The values are averaged across subjects (± SEM).

Among the models tested, the UGM provided the most comprehensive fit. Although its overall performance was comparable to that from the DDM-collapsing - with no significant difference in the BIC values of each model (paired-samples Wilcoxon signed-rank test, FDR-corrected for multiple comparisons, p > 0.01) – the UGM was the only model able to capture the observed differences in DTs across all trial types (Fig 5D-F; paired-samples Wilcoxon signed-rank test, z = 3.70 – 4.01, FDR-corrected for multiple comparisons, p < 0.001). Additionally, the UGM accurately reflected the lack of significant differences in the DT modulation between bias- for and bias-against trials when comparing no-pause and pause conditions (paired-samples Wilcoxon signed-rank test, p > 0.95).

Despite its strength, the UGM, like the other models, fell short in explaining the effects of temporal disruptions introduced by a gap. While the DDM-fixed and UGM both captured overall differences in DTs between no-pause and pause conditions (Fig 5G-I-left), the DDM- collapsing failed to do so. All models correctly replicated the observed difference in the number of tokens used for decisions between pause and no-pause conditions (Fig 5G-I-middle). However, none of the models, including the UGM, were able to explain the lack of a significant difference in accuracy between no-pause and pause conditions (Fig G-I-right).

Overall, although the UGM outperformed the other models in capturing key aspects of the data, its inability to explain the preservation of accuracy in pause trials highlights a crucial limitation. This limitation raises an important question: Are we overlooking essential aspects of how pieces of information are combined during decision making? Specifically, could the temporal dynamics of the trial – such as the sequence and timing of evidence presentation – play a pivotal role in shaping behavior?

### Perceptual evidence is weighted differently

To investigate whether the contribution of each token was influenced by its temporal position within a sequence, we used logistic regression (see Materials and Methods). This analysis allowed us to compute regression coefficients that estimate the weight assigned to each token in the decision-making process. Moreover, we could also estimate the potential neuronal noise (signal-to-noise ratio, SNR) as the average weight assigned to all tokens, and potential side biases (Keung et al., 2020).

First, we computed the estimated weights for all experimental trials, considering decisions made within the first 12 tokens, as most decisions were made in that range (99.27 ± 0.23%, ± SEM). The SNR was significantly greater than zero (one-sample Wilcoxon signed-rank test, z = 4.01 and Cohen’s d=2.87, FDR-corrected for multiple comparisons p <0.001; Fig 6A- left), indicating that participants based their decisions on the evidence provided by the tokens. To assess whether participants weighted all tokens equally, we subtracted the average weight from the individual weight of each token. This analysis revealed that participants did not assign equal weight to all tokens, but rather, they weighted tokens unevenly over time. Specifically, tokens presented towards the middle of the sequence had a stronger impact on decisions, while tokens presented later in the sequence, particularly those towards the end, were given significantly less weight (one-sample Wilcoxon signed-rank test, z = 3.46 – 4.01, FDR-corrected for multiple comparisons, p < 0.001; Fig 6A-middle). Importantly, there was no side bias in the choices (Fig 6A-right).

**Figure 6.**
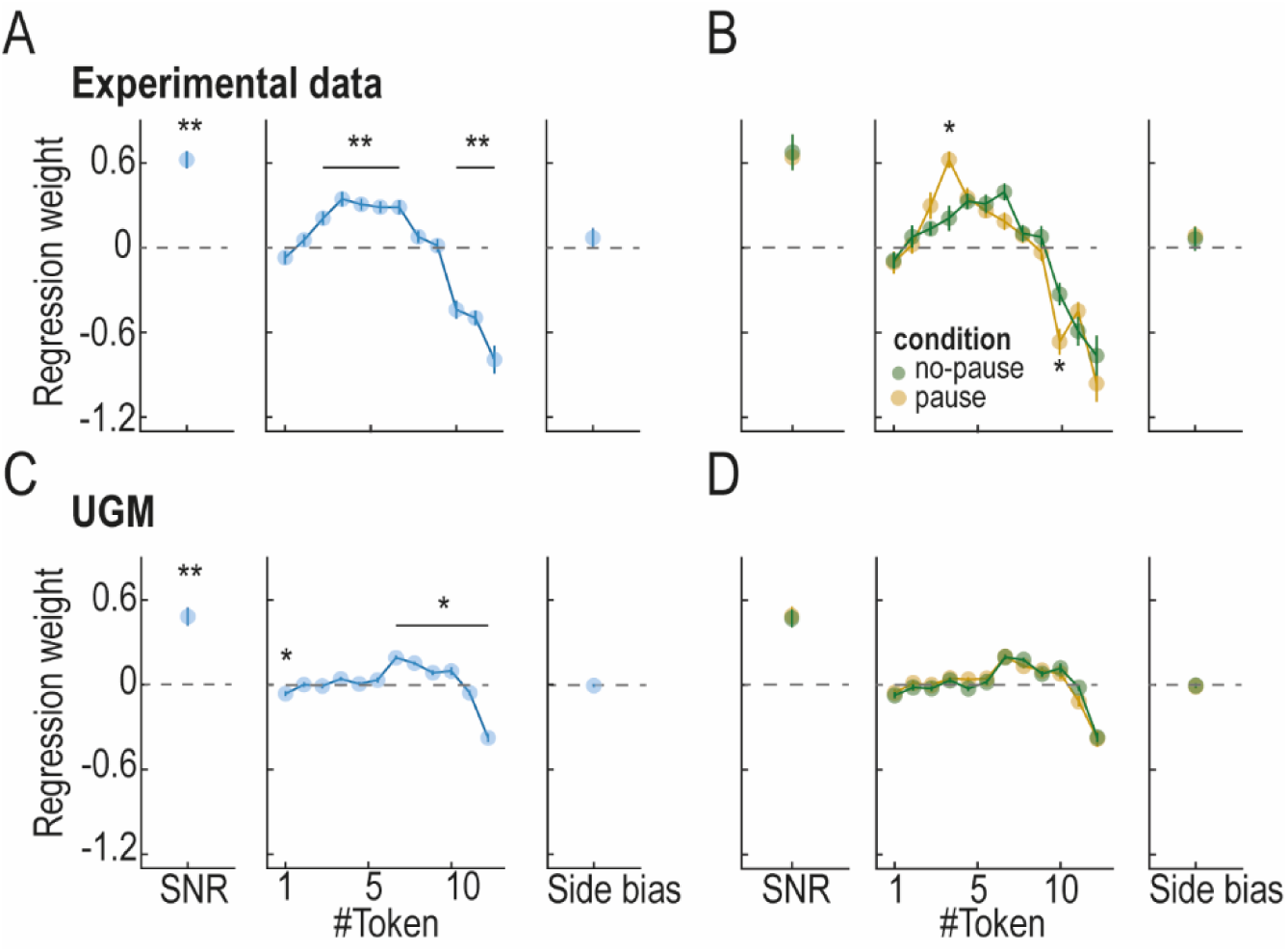
Integration kernels for real and simulated data. **(A)** SNR (*left*), deviation of tokens weight from the mean (*middle*) and side bias (*right*) for all experimental trials (one-sample Wilcoxon signed-rank test, false discovery rate (FDR)-corrected for multiple comparisons, ** = p <0.001). **(B)** SNR (*Left panel*), deviation of tokens weight from the mean (*middle*) and side bias (*right*) for experimental data, divided into no-pause and pause conditions (paired-samples Wilcoxon signed-rank test, false discovery rate (FDR)-corrected for multiple comparisons, * = p <0.05). **(C-D)** Same conventions as in (A) and (B) for simulated data obtained from the UGM (* = p <0.01).

Next, we examined whether this uneven weighting was consistent across no-pause and pause conditions. The proportion of trials with decisions made within the first 12 tokens did not significantly differ between no-pause and pause trials (99.20±0.23 and 99.33±0.14, ±SEM, respectively; paired-samples Wilcoxon signed-rank test, p > 0.58). No significant differences were found in SNR or side bias between conditions (Fig 6B-left and Fig 6B-right). However, the most striking result emerged for the 4^th^ token, which occurred immediately after the pause. This token was weighted significantly more strongly in the pause condition than in the no-pause condition (paired-samples Wilcoxon signed-rank test, z = 3.53, FDR-corrected for multiple comparisons, p < 0.01; Fig 6B-middle). This suggests that the token presented right after the pause has a disproportionately strong influence on decision making in the pause condition, highlighting the critical role of the temporal gap in shaping how information is integrated during the decision process. Notably, the three first tokens, presented before the temporal gap in the pause condition, were weighted similarly to those in the no-pause condition, supporting the notion that the pause does not degrade the information provided earlier in the sequence.

We performed the same computations with the simulations obtained with the UGM. We found that the SNR was significantly above zero (one-sample Wilcoxon signed-rank test, z = 4.01 and Cohen’s d=2.16, FDR-corrected for multiple comparisons, p <0.001; Fig 6C-left), indicating that the model assigned meaningful weights to the tokens. However, unlike the real data, where late tokens were weighted less, the simulations show a slight increase in weight for later tokens (one-sample Wilcoxon signed-rank test, z = 2.73 – 4.01, FDR-corrected for multiple comparisons, p < 0.05; Fig 6C-middle). This divergence highlights the model’s inability to replicate the observed temporal patterns of tokens weighting accurately. Additionally, no side bias was observed in the simulated data (Fig 6C-right), and comparisons between no-pause and pause conditions revealed no significant differences in either SNR (Fig 6D-left) or side bias (Fig 6D- right). Crucially, unlike the experimental data, where the token presented immediately after the pause (the 4^th^ token) received disproportionally higher weight in pause trials, the simulations showed no significant differences in the weighting of this token between conditions (paired-samples Wilcoxon signed-rank test, p > 0.05; Fig 6D-middle).

Overall, while the simulations captured some general features of the data, they failed to replicate key experimental results. In particular, the shift in tokens weighting and the absence of enhanced weighting of the post-gap token highlight the model’s limitation. These discrepancies underscore the need for additional mechanisms to explain behavior under these conditions. Such mechanisms may involve how the brain dynamically integrates evidence during decision making or how it processes the temporal disruption introduced by the pause.

Importantly, the question of why accuracy remains consistent between no-pause and pause conditions persists. Could the inherent statistical properties of the perceptual stimuli make the post-gap token highly informative? If so, assigning greater weight to this token may compensate for the expected reduction in performance in the pause condition. A question we address next.

### Stimulus statistics enhance accuracy in the pause condition

To assess the contribution of each token to the correct option, we calculated the proportion of trials in which each specific token favored the correct target. In most cases, the majority of tokens favored the correct choice (Fig 7A-left). We then used these favorable cases to compute the information added to the SP at the time of each token’s jump (see Materials and Methods). Notably, due to the structured nature of the task, where specific profiles of token sequences were used, the 4^th^ token emerged as one of the most salient pieces of information (Fig 7A-right). Choices were significantly biased by the direction of that token more often in the pause than in the no-pause condition (paired-samples Wilcoxon signed-rank test, FDR-corrected for multiple comparisons, z = 3.49, p < 0.01; Fig 7B-left), consistent with the higher weight given to the token for the pause condition. This bias resulted in a significant improvement in accuracy (paired-samples Wilcoxon signed-rank test, z = 3.53, FDR-corrected for multiple comparisons, p < 0.01; Fig 7B-right), indicating that the 4^th^ token played a crucial role in enhancing performance. We hypothesize that this is the reason for the maintenance of similar accuracy levels between the two conditions.

**Figure 7.**
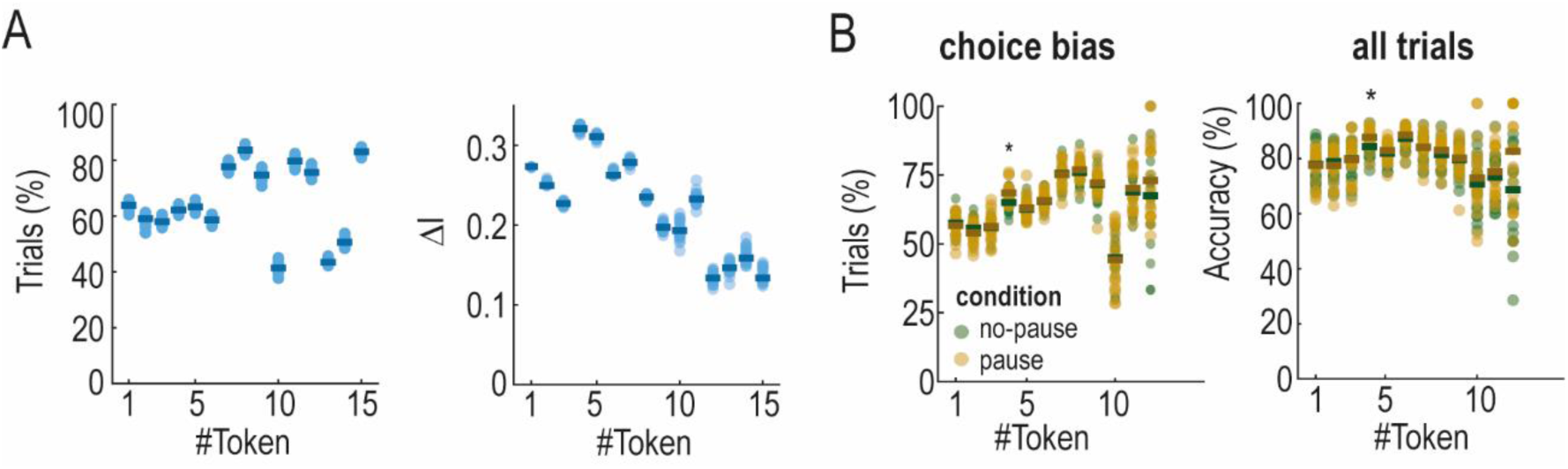
Performance by direction of tokens jump. **(A)** Proportion of trials in which the jump of the specific token favored the correct target (*left*). Variations in information added by each specific token (*right*). **(B)** Proportion of choices made towards the target that received the jump of the specific token (*left*). Accuracy for trials in which the specific token jumped towards the correct target (*right*; paired Wilcoxon signed rank test, FDR-corrected for multiple comparisons, * = p<0.05).

To confirm this, we designed a non-structured task in which there were no specific trial profiles. Instead, the side of each token’s jump was randomly selected from a Bernoulli distribution with mean 0.71 for the first 12 tokens (see Materials and Methods). As a result, the jumps of these tokens consistently favored the correct target in a proportion around that mean (Fig 8A-left). Unlike the structured task, the information provided by the 4^th^ token was no longer salient, as the information added by each token gradually diminished over time (Fig 8A-right). In this case, DTs and the number of tokens at decision followed a similar trend than in the non-structured task (paired-samples Wilcoxon signed-rank test, z = 5.03 – 5.09, p < 0.0001), with an increase and decrease, respectively, between no-pause and pause conditions (Fig 8B).

**Figure 8.**
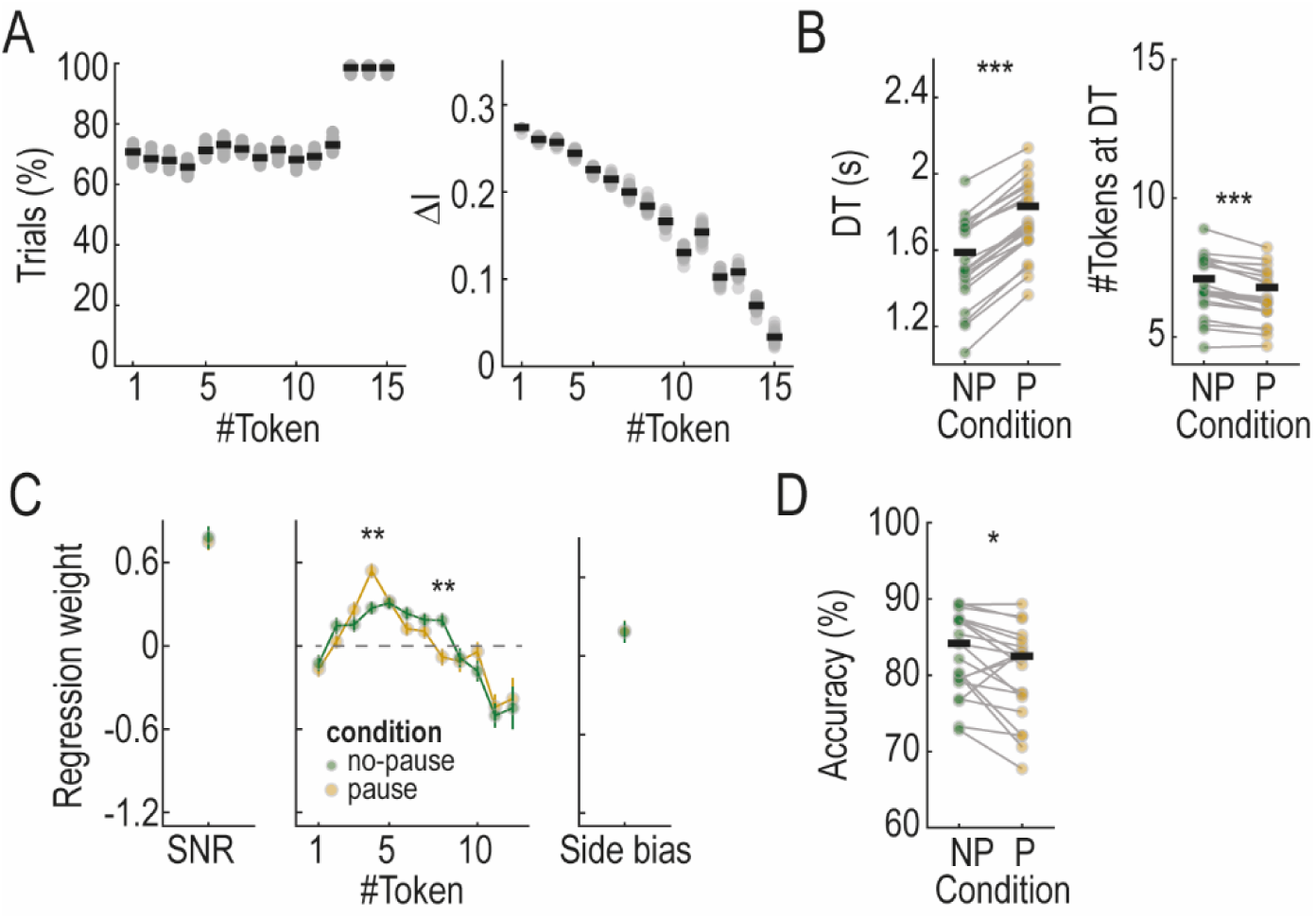
Stimulus statistics, behavioral outcome and information weight in the non- structured task. **(A)** Proportion of trials in which the specific token favors the correct target (*left*) and information added by each token’s jump (*right*). **(B)** DTs and number of tokens at decision time for no-pause (NP) and pause (P) conditions. **(C)** SNR *(left*), deviation of tokens weight from the mean (*middle*) and side bias (*right*) for trials divided in no-pause and pause conditions (*** = p<0.001). **(D)** Accuracy in no-pause (NP) and pause (P) conditions (paired Wilcoxon signed rank test, FDR-corrected for multiple comparisons, * = p<0.05, *** = p<0.0001).

Despite the absence of a structured sequence and the lack of saliency of the 4^th^ token, the weight assigned to this token remained significantly greater in the pause condition compared to the no-pause condition (Fig 8C-middle). This replication underscores that the enhanced impact of the information following a temporal gap is a robust phenomenon, independent of the saliency of the event itself. Importantly, the SNR and side bias effects did not differ between conditions (Fig 8C-left and 8C-right). However, unlike the structured task, accuracy was significantly lower in the pause condition (paired-samples Wilcoxon signed-rank test, z =3.10, p < 0.01), supporting our hypothesis that accuracy was preserved in the structured task due to the saliency of the post-gap token. These findings provide compelling evidence that the brain assigns disproportionate weight to information arriving immediately after a temporal gap, even when that information lacks inherent saliency.

### Evidence weighting is shaped by the temporal gap

A potential confound in our results on the differential weighting of information between no-pause and pause conditions lies in the difference in the number of tokens at the time of decision. Specifically, the disproportionate weight assigned to the post-gap token in the pause condition could reflect decisions being made with fewer tokens and, consequently, earlier in the tokens sequence. To address this, we conducted an additional analysis where we calculated the integration kernel for no-pause and pause conditions after applying a bootstrapping procedure to equalize the distribution of the numbers of tokens at decision across conditions (see Materials and Methods). This procedure was applied for both the structured and non-structured tasks, resulting in identical numbers of tokens at decisions for the no-pause and pause conditions in each task (mean ± SEM: 8.24 ± 0.15, 7 ± 0.21, for the structured and non-structured tasks, respectively). Consistent with our previous results, the 4^th^ token received significantly greater weight in the pause compared to the no-pause condition in the two tasks (Fig 9; paired Wilcoxon signed rank test, z = 2.90 – 2.99, FDR-corrected for multiple comparisons, p < 0.05). These results confirm that the differential weighting of evidence between conditions is indeed associated with the temporal gap. This finding suggests the involvement of cognitive mechanisms specifically tuned to process and prioritize information following interruptions, a phenomenon that traditional decision-making models fail to capture.

**Figure 9.**
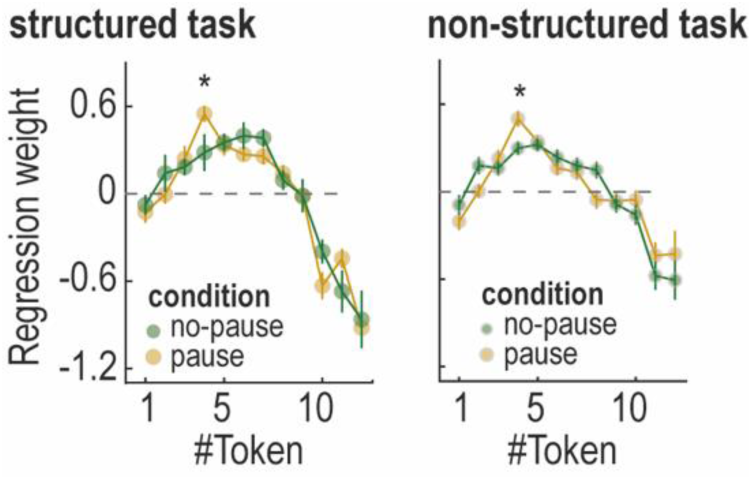
Information weighting is related to the temporal gap. Deviation of tokens weight from the mean in the structured (*left*) and non-structured tasks (*right*) when a bootstrapping procedure was conducted to ensure an equalized number of tokens at decision for no-pause and pause conditions (paired Wilcoxon signed rank test, FDR-corrected for multiple comparisons, * = p < 0.05).

## Discussion

Our findings reveal that a temporal gap, during which no information is presented, significantly influences how information is processed for decision making. Specifically, we demonstrate that when a temporal gap occurs within a sequence of perceptual stimuli, decisions are made with less information compared to sequences without a gap. Additionally, the temporal gap alters the way individual pieces of information are weighted, with post-gap information exerting a disproportionately strong influence on choices. Despite this reduction in available information, accuracy remained stable in a structured task where the post-gap token was particularly informative due to the statistical properties of the stimulus. To test whether the increased weighting of post-gap information persisted regardless of stimulus structure, we designed a non-structured task in which the saliency of the post-gap token was reduced. While accuracy was no longer preserved, the post-gap token still carried disproportionate weight, indicating that its influence is independent of its intrinsic saliency.

Traditional decision-making models failed to replicate key aspects of our findings, including the preservation of accuracy in the structured task and the uneven weighting of sequential information – particularly the heightened influence of the post-gap token. This limitation suggests that existing models may lack critical mechanisms necessary to fully capture behavior under these conditions. Our results highlight the need for decision-making models to incorporate not only stimulus statistics and dynamic evidence integration but also mechanisms that account for the increased salience of information following temporal disruptions.

### Task design and generalization

The position and duration of the temporal gap were carefully selected to maximize its potential impact on decision making. On the one hand, the gap was placed at a point in the sequence that allowed us to test whether disrupting early information would alter its influence on choices. This was assessed by comparing bias-for and bias-against trial types. If the gap weakened the contribution of early evidence, we would expect a differential effect between these trial types in the pause versus no-pause conditions. Additionally, the gap was placed at a point in the sequence that ensured its presentation on the majority of trials, as decisions were typically made later in the sequence. On the other hand, the gap’s duration was chosen to introduce a meaningful disruption by breaking the symmetry of the sequence with a duration that differed from the token jumps. Moreover, it was selected to be long enough to exert an effect on behavior, but short enough to avoid conscious detection, as confirmed with a questionnaire. An important open question is whether variations in the gap’s position or duration would lead to different effects on the decision-making process.

Our task used on non-noisy sensory information, raising the question of whether these findings extend to scenarios involving noisy stimuli (Roitman and Shadlen, 2002; Kiani et al., 2013; Marcos et al., 2015). However, a previous study using a task similar to ours but with noisy stimuli revealed no discernable difference in decision-making dynamics compared to those observed with non-noisy stimuli (Cisek et al., 2009; Thura et al., 2012; Ferrucci et al., 2021). Given these findings, it is reasonable to infer that temporal gaps would exert similar effects in both noisy and non-noisy environments. Nevertheless, future experiments should explicitly test this hypothesis by introducing temporal gaps in sequences of noisy information.

### Temporal disruptions and decision-making models

In recent decades, various decision-making theories have emerged. The prevailing view suggests that decisions are made by accumulating sensory evidence until it reaches a decision boundary (Stone, 1960; Van Zandt et al., 2000; Brown and Heathcote, 2008; Churchland et al., 2011; Bollimunta et al., 2012; Cassey et al., 2014; Dutilh et al., 2019). Although this framework could explain many behavioral and neuronal observations, it fails to explain behavior when information continuously fluctuates (Cisek et al., 2009; Winkel et al., 2014; Holmes et al., 2016; Ferrucci et al., 2021; Trueblood et al., 2021), as it commonly occurs in natural conditions. In such cases, models incorporating an urgency signal – a time-varying drive to commit to a choice – better fit experimental data (Cisek et al., 2009; Carland et al., 2016; Ferrucci et al., 2021). Urgency-related neural dynamics have been observed in regions such as the dorsal premotor and primary motor cortex, as well as the basal ganglia (Thura and Cisek, 2014, 2017). However, urgency may influence decision making either by modulating sensory evidence or by collapsing decision bounds over time (Ditterich, 2006b, a; Churchland et al., 2008; Cisek et al., 2009; Hawkins et al., 2015; Carland et al., 2016; Murphy et al., 2016; Ferrucci et al., 2021). To examine these models under conditions where temporal gaps interrupted sensory evidence, we tested three implementations: a DDM (without urgency), a DDM-collapsing (urgency as collapsing decision bounds), and a UGM (urgency as a time-increasing signal). Although prior studies showed that the DDM fails in fluctuating environments (Cisek et al., 2009; Ferrucci et al., 2021), we included it to assess whether it could still capture decision dynamics under temporal disruptions. However, it failed to do so. Among the urgency-based models, the UGM performed best, reinforcing the idea that urgency is essential in dynamic environments. However, it still could not explain the disproportionate weighting of the post-gap stimulus, suggesting the involvement of additional underlying mechanisms. One possibility is that temporal gaps create a contrast effect, making the post-gap stimulus stand out more prominently and exert greater influence on decisions. This highlights the need for further refinements to current models to fully capture how the brain integrates information over time.

In scenarios characterized by intermittent information, decisions cannot be made instantaneously, necessitating the use of memory to construct and maintain a representation of absent information while integrating new inputs as they become available. This dual demand requires memory systems to strike a delicate balance: they must be stable enough to preserve information over time, yet sufficiently flexible to adapt when new information arrives. In this study, we have proposed a basic equation to model this process, capturing how previous information is retained but gradually degrades as new information is incorporated. However, the neural mechanisms underlying this computation within distributed memory networks remain unresolved. One compelling hypothesis is that a population of neurons with heterogeneous excitability interact with each other, giving rise to stable yet adaptable dynamics (Marcos et al., 2019). Nevertheless, to provide a definitive answer to this question, further experiments involving animals and electrophysiology are imperative.

Our results demonstrate that temporal gaps significantly modulate decision making, reducing the amount of evidence needed for a choice and altering how evidence is processed. These results challenge the idea that behavior remains invariant when delays are present (Kiani et al., 2013; Waskom and Kiani, 2018). Both experimental and computational analyses indicate that the proposed “freezing” mechanism, which suggests stability of the decision-making process during gaps, is insufficient to explain the observed behavior. Additionally, a pipeline processing of perceptual information (Moran et al., 2015; Calder-Travis et al., 2023), which accounts for slow integration of sensory information, also fails to capture the dynamics of decision making during temporal gaps. Together, these findings underscore the need for more refined models that can capture the complex interplay between temporal disruptions and evidence processing in decision making.

### Weighting of dynamic evidence

Previous research has shown that when participants are presented with two pulses of perceptual stimuli, the second pulse exerts a stronger influence on decisions (Kiani et al., 2013). While these studies have provided valuable insights, they have not fully explored how individual pieces of information shape decision making over time. By using stimuli that changed in discrete steps, our study allowed for the precise isolation and quantification of the contribution of each piece of information to decision making. We revealed an uneven weighting of information across the sequence, with this weighting being further modulated by the information immediately following a temporal pause. These findings highlight the critical role of temporal context in shaping perceptual decisions.

Similar patterns of uneven weighting of information haven been reported in prior studies (Keung et al., 2019, 2020), where participants were presented with sequences of auditory “clicks”, and asked to determine which side (left or right ear) received the most clicks. Integration kernels in these studies varied in shape, with the most common being a “bump” shape where information in the middle of the sequence was weighted more heavily. The authors proposed a divisive normalization model that successfully captured the variability in kernel shapes and the observed behaviors (Keung et al., 2020). However, applying this model to our data presents several challenges. Notably, the model does not account for the time between events, treating them as continuously occurring with instantaneous durations. However, in our experimental design time is critical: while the tokens are presented at consistent intervals, the temporal gap introduces an asymmetry by extending the sequence duration. This temporal disruption is integral to our study and influences the decision-making process in a way that the divisive model cannot currently address. Additionally, in this model, decisions are made at the end of the sequence presentation, by comparing the activity of two population units. This approach does not involve an explicit decision threshold, and the choice is based on which population exhibits higher activity at the end of the sequence. Incorporating these features to the model would imply significant theoretical adjustments, introducing complexities and implications that fall beyond the scope of our study.

Importantly, our study differs from previous work that found invariant decision making in the presence of temporal gaps (Kiani et al., 2013; Waskom and Kiani, 2018; Tohidi-Moghaddam et al., 2019; Azizi and Ebrahimpour, 2023). A key distinction lies in the experimental design: while those studies used tasks with a go signal instructing participants when to decide, our task allowed participants to respond at their discretion. This design potentially enabled them to manage their speed-accuracy trade-off through an urgency signal that grows over time (Cisek et al., 2009). Additionally, the perceptual evidence in our task was dynamic, changing in discrete steps throughout the trial. These differences in experimental designs highlight the importance of considering both accuracy and decision times as complementary behavioral measures for a comprehensive understanding of the decision-making process. Moreover, incorporating time-varying information into the experimental designs – reflecting real-world scenarios – is essential for advancing our understanding of perceptual decision making in naturalistic contexts.

### Conclusions

Overall, our findings emphasize the importance of investigating decision making with sensory stimuli that reflect real-world dynamics to gain deeper insights into perceptual decision making. By isolating the contribution of each piece of evidence, we observed an uneven weighting, particularly influenced by the gaps between stimuli. This highlights the need for decision-making models that can better capture these dynamics and the complexities of how information is processed across time. Refining computational models to account for temporal disruptions, along with incorporating time-varying information into experimental designs, will be crucial for advancing our understanding of perceptual decision making in naturalistic environments.

## Materials and Methods

### Ethics statement

All experimental procedures were in accordance with the ethical standards of the university research committee, with the Code of Ethics of the World Medical Association (Declaration of Helsinki, 1964) and its later amendments. The experimental protocol was approved by the Ethics Committee of the Miguel Hernández University/CSIC/Institute of Neurosciences of Alicante. Before performing the tasks, all participants provided written consent.

### Experimental tasks

Each participant completed one of two experimental tasks: the structured or the non-structured task. Within each task, participants were required to complete a set number of blocks, each with a predefined number of correct trials, to prevent random guessing. The experimental paradigm was based on the modified version of the task from Cisek et al. (2009), as presented in Ferrucci et al. (2021).

In the two tasks, at the beginning of each trial, three circles with white outlines were displayed on a black screen, each measuring 2.5 cm in diameter (Fig 1A). The central circle, positioned at the center of the screen, contained 15 tokens randomly distributed within its area. Two additional circles, designated as targets, were placed 5 cm on either side of the central circle (left or right). To initiate a trial, the participant was required to place the mouse cursor inside the central circle. Subsequently, the tokens began to jump to one of the two peripheral circles every 200 ms, disappearing just before the subsequent token’s jump. Participants were free to report their decision at any moment before the last token’s jump, and were instructed to report their choice by moving the cursor inside one of the peripheral circles. Following their selection, the outline of the chosen circle changed from white to green for correct choices and to red for incorrect choices. After feedback, the remaining tokens jumped to one of the peripheral circles every 20 ms. Consequently, participants could save time by making rapid choices, encouraging the participants to respond before the trial concluded. If the choice was not made before the last token’s jump, no feedback was provided. A time interval of 500 ms separated the end of a trial from the beginning of the next.

Two conditions were introduced in the sequence of jumps: trials with a 300 ms temporal gap before the jump of the fourth token, termed *pause trials*, and trials without such a gap, termed *no-pause trials*. The temporal gap consisted of a period where no jumps occurred and no information was available on the screen. The specific position of the temporal gap within the token’s sequence was selected to maximize its potential effect on behavior (see below). Within each block, half of the trials were pause trials and both conditions were randomly interleaved within the block.

### Structured task

Twenty-one participants performed the structured task (aged 23-42; all right-handed except one; ten females). Each participant performed 5 blocks of perceptual decision-making trials that contained pause and no-pause trials in a random order. The participant had to make 100 correct trials to complete each block. A BENQ GL2580 monitor (24.5’’) was used to display the visual stimuli for the task and a mouse was used as an interface between the participants and the computer. The participants sat in front of the screen at approximately 60 cm.

In each of the five blocks, we included five trial types, in equal proportion (20%): “random”, “easy”, “ambiguous”, “bias-for” and “bias-against” (Cisek et al., 2009; Ferrucci et al., 2021). In random trials, the direction of token jumps was randomly determined. Easy trials involved most tokens jumping to the correct target while ambiguous trials featured an even distribution of tokens between the two targets until just before the trial’s end (Fig 1C-left). The bias-for and bias-against trials were particularly effective in discerning the effect of the temporal gap (Ferrucci et al., 2021). The reason is that these two trial types differ solely in the direction of the first six tokens’ jumps, which is completely opposite, with the highest divergence occurring after the 3^rd^ token jump (Fig 1C-right). By using these well-defined sequences, the analysis can specifically target how the temporal gap influences the processing of early vs late information. Introducing a temporal gap immediately after this jump has to potential to substantially impact the behavioral differences between the two trial types by altering the processing of early versus late information. Random trials were included to prevent subjects from anticipating the trial pattern.

### Non-structured task

Thirty-four participants performed the non-structured task (aged 18-48; all right handed except one; twenty-two females). Each participant completed 6 blocks with no-pause and pause trials randomly interleaved, and was required to perform 125 correct trials. A laptop (Lenovo IDEAPAD 310-15ABR; screen: 15.6’’, resolution: 1920x1080) was used to display the visual stimuli, and a mouse served as the interface between the participants and the computer. The participants sat approximately 60 cm from the screen.

For each trial, the jump of the first 12 tokens was generated following a Bernoulli distribution with a mean probability of 0.71 for landing on the correct target. The last 3 tokens always jumped towards the correct target.

### Behavioral analysis

At each time in the trial, we could calculate the “success probability” *p*_*i*_(*t*) associated with being correct when selecting target *i*. This probability was determined based on the number of tokens remaining in the center (*Nr*), the number of tokens at target 1 (*N1*) and the number of tokens at target 2 (*N2*). Specifically, the probability of target 1 being correct was computed as follows (Cisek et al., 2009; Ferrucci et al., 2021):

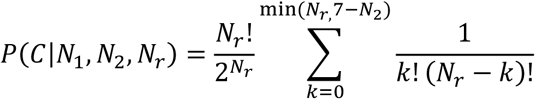

The SP was calculated by determining the probability that the chosen target was correct.

To estimate decision times (DTs), participants performed an additional set of 40 trials where only one token jumped to one of the targets, randomly. The interval from the token’s arrival at the target to the moment the mouse cursor left the central circle to report the choice established the subject’s baseline reaction time (RT). The baseline RT was subtracted from the RT recorded from the main task. The RT in the main task was estimated by subtracting the trial’s start time from the time of maximum acceleration of the cursor. By employing this method, we mitigated the influence of trials in which participants moved the cursor from the central circle but experienced delays in making their selection (∼10% of trials). We estimated a mean RT of 0.32 s (± 0.01, SEM) for the structured task and a mean RT of 0.34 s (± 0.01, SEM) for the non-structured task. Number of tokens and SPs were then computed based on the estimated DT.

We computed the amount of information added by each token (Δ*I*) by calculating the SP value when the token appears (*SPtoken*) and the one just before it (*SPpre-token*). Then, we calculated the logarithmic difference between the two:

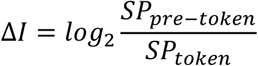

Unless otherwise stated, all analyses are performed on correct trials.

We controlled the lack of awareness of either the temporal gap or the trial types with a questionnaire, provided at the end of the experimental session. Only one participant in each task reported the presence of the gap.

### Computational model

We fitted the experimental data from the structured task to three different decision-making models. All models received the same sensory evidence input, which was derived from a module simulating the degradation of perceptual information due to retention in memory.

### Sensory evidence

The perceptual evidence was updated with each new token jump, simultaneously adding new information while causing the retained evidence to decay:

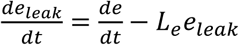

where *Le* represents a leakage term for the retained perceptual information, and 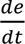is the change in the perceptual evidence displayed on the screen. Specifically, 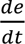 is defined as:

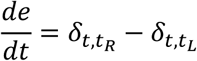

where the delta functions represent the jump of a token to either right (*δ*_*t*,*tR*_ = 1) or left (*δ*_*t*,*tL*_ = 1). Since the tokens’ jumps are deterministic, we modeled the perceptual evidence as a deterministic process as well.

### Decision-making models

We implemented the drift diffusion model with fixed boundaries (DDM-fixed), the drift diffusion model with collapsing boundaries (DDM-collapsing) and the urgency-gating model (UGM). The models have been selected based on the validity of the three to explain a wide range of experimental data (Stone, 1960; Laming, 1968; Ratcliff, 1978; Usher and McClelland, 2001; Mazurek et al., 2003; Bogacz and Gurney, 2007; Hawkins et al., 2015; Ferrucci et al., 2021; Yau et al., 2021; Smith and Ratcliff, 2022).

The basic idea behind the diffusion process is that evidence accumulates towards one of two decision bounds, and a decision is made when the accumulated evidence reaches a bound. The predicted response time is the sum of the time it takes for the process to reach a bound and the additional time required for non-decision-related components of choice, including action execution. The dynamics of this process are governed by five key parameters: the average rate of drift towards one bound (drift rate), the noise in the diffusion process, the separation of the bounds, the starting position of the diffusion process and the time taken for non-decision-related processes (Stone, 1960). Over recent decades, variability has been introduced into some of these parameters, such as drift rate.

In our implementation of the process, we modeled the diffusion as starting at 0 and drifting toward two symmetric bounds and estimated DTs as the time the process takes to reach one of the two bounds. Non-decision-related times were not included, as we aimed at fitting DTs estimated from the experimental data. We accounted for variability in the drift diffusion rate, which varied from trial to trial, and modeled the process as stochastic. Therefore, the DDM-fixed and DDM-collapsing followed these same dynamics:

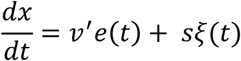

where *x* describes the state of the process at time *t* (decision variables), *v*′ is the drift rate, *e* is the sensory evidence, *s* is a scaling factor and *ξ* is an independent and identically distributed random sample taken from a standard normal distribution. We used the Euler’s method (Brown et al.) to simulate the model:

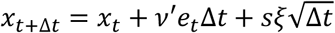

where Δ*t* is the step size of the simulation. In each simulated trial, the initial value of *x* was set to 0 and the drift rate ν’ was sampled from a normal distribution with standard deviation *η* and mean*ν*. A decision was considered to be made when *x_t+Δ*t*_* reached a specific value *θ* (bound), and the decision time was estimated as 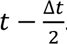. In the DDM-fixed, *θ* was kept constant throughout the trial, whereas in the DDM-collapsing *θ* varied according to:

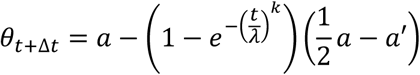

where λ is the scale parameter of the Weibull distribution, *a*′ defines the extent to which the bounds collapsed, and *k* determines the shape of the bound.

In the simulations, we arbitrarily set *s* to 0.1 and Δ*t* to 0.01 s. The value of *k* was set to 3 in the DDM-collapsing to impose a “late collapse” of the bounds, because it is a representative value when the value is freely estimated (Hawkins et al., 2015). The parameters *ν*, *η* and *θ* in the DDM-fixed were free parameters estimated with a fitting procedure optimized for the goodness of fit, whereas *ν*, *η, a*, *a*′ and λ were estimated for the DDM-collapsing, following the same fitting procedure. In both cases, the leaky term in the sensory evidence (*L*_*e*_) was also fitted.

In the urgency-gating models, the build-up activity towards a bound is driven by a time-varying gain signal rather than by temporal accumulation of evidence. In these models, momentary evidence is combined with a growing signal that reflects increasing urgency to make a decision. The UGM followed the stochastic equation:

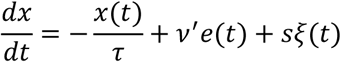

where *x*, *v*’, *e*, *s*, ξ are parameters defined as in the DDMs and *τ* is the time constant of a low-pass filter. We used the Euler’s method (Brown et al., 2006) to simulate the model:

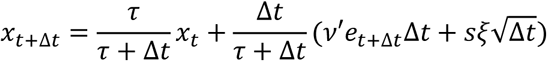

where Δ*t* is the step size of the simulation. We set *s* and Δ*t* as in the DDMs and τ to 0.01 s. An urgency signal, which increases with the time elapsed since the initiation of the decision-making process (*ut=t*), multiplied the instantaneous value of the accumulation of evidence. The decision was considered to be made when *x*_*t*+Δ*t*_ · *u*_*t*+Δ*t*_ > *θ* or *x*_*t*+Δ*t*_ · *u*_*t*+Δ*t*_ < −*θ* and the corresponding decision time was estimated as 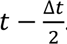. The parameters *ν*, *η, θ* and the leaky term *L*_e_ were estimated following the same procedure as for the DDMs.

### Estimation of model parameters

The free model parameters of the models were estimated independently for each model to fit the participants’ data. We used the quantile maximum products estimation (QMPE) (Heathcote et al., 2002; Ferrucci et al., 2021), dividing DTs into quantiles for correct and error trials. The QMPE estimates the similarity between simulated and real data by comparing the proportion of data within each quantile. We used a differential evolution algorithm to maximize the goodness of fit (Mullen et al., 2011; Ardia et al., 2013), with broad parameters boundaries and executing 100 particles for 500 search iterations. To mitigate the risk of local minima, this parameter estimation process was repeated five times for each model and participant’s data. Model predictions were assessed through Monte Carlo stimulation, generating 10,000 replicates for each experimental condition. Upon termination of the search, data was simulated using the parameters set that yielded the highest goodness of fit.

The data fitting procedure was performed using all trials, including correct and errors, across pause and no-pause conditions.

### Integration kernel

To calculate the contribution of each specific token to the choice made, we used logistic regression (Keung et al., 2020). In this approach, the probability of choosing “left” was estimated as:

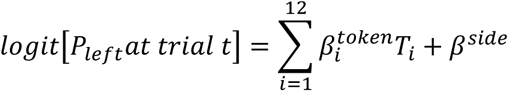

where Ti represents the *i*-th token (+1 for left, -1 for right and 0 if the tokens was not presented). The regression coefficients (*β*^*token*^_*i*_, *β*^*side*^) reflect the contribution of each factor: the perceptual tokens and side bias, respectively, on the decision. To fit the model, we considered the first 12 tokens because participants made most of their decisions within that range (99.27 ± 0.23% and 99.59 ± 0.17%, for the structured and non-structured tasks, respectively, ± SEM). We estimated the signal-to-noise ratio (SNR) as the average weight assigned to all tokens. Correct and error trials were considered.

For the analysis with the data sorted by no-pause and pause conditions, only trials with decisions made after the 4^th^ token jump were included.

We used the same procedure to estimate the model-predicted kernel and SNR.

### Equalization of the number of tokens at decision

To equalize the number of tokens at the time of decision between no-pause and pause conditions, we implemented a bootstrapping procedure. For each token position (from token 4 to token 12), we selected the trials where decisions were made at that specific token in both the no-pause and pause condition, separately. For each token, we determined the smaller number of trials to sample randomly from each condition. This procedure was repeated 1,000 times to ensure robustness.

### Statistical tests

We used a non-parametric test for statistical comparisons. Specifically, the Wilcoxon signed-rank test was employed, as it does not assume normality in the data distribution. Moreover, it is less sensitive to outliers, providing a reliable measurement of central tendency by focusing on the median rather than the mean.

Correction for multiple comparisons was performed using the False Discovery Rate (FDR), to account for Type I errors.

## Data and code availability

The data and code for computational modelling used in this study, as well as any additional information are available from the corresponding author upon request.

## Acknowledgements

The authors are grateful to Paul Cisek for discussion and comments on the manuscript. This research was financially supported by Grant RYC2021-035061-I and Grant PID2022- 141173NA-I00, funded by MICIU/AEI/10.13039/501100011033 and by the European Union NextGenerationEU/PRTR and ERDF/EU, respectively, awarded to EM; by Grant PID2021- 128158NB-C21, funded by MICIU/AEI/10.13039/501100011033 and ERDF/EU, awarded to SC; and by the Programs for Centres of Excellence in R&D Severo Ochoa, Agencia Estatal de Investigación (CEX2021-001165-S)

